# Reduced susceptibility to experimentally-induced complex visual hallucinations with age

**DOI:** 10.1101/2025.05.27.656359

**Authors:** Oris Shenyan, Laura Haye, Georgia Milne, Matteo Lisi, John A. Greenwood, Jeremy Skipper, Tessa M. Dekker

**Affiliations:** Experimental Psychology, Division of Psychology and Language Sciences, University College London, London, UK; Institute of Ophthalmology, University College London, London, UK; Department of Psychology, Royal Holloway University, London, UK

## Abstract

Visual hallucinations occur across many clinical conditions, but can also be induced experimentally in healthy individuals, using high-frequency flicker (Ganzflicker) and sensory deprivation (Ganzfeld). It is unclear how hallucinatory proneness changes across the lifespan, with prior questionnaire-based studies showing mixed results. As factors such as multi-sensory acuity loss and relatively increased reliance on prior knowledge may increase as we age, and these are considered risk factors for hallucination proneness, we hypothesised that reported decreases in hallucinations might reflect stigma-related underreporting. We therefore sought to measure hallucination proneness in 44 healthy adults spanning the adult lifespan (younger age group; n=22; age 19-39 years, mean 27.2 ± 5.5; older age group n=22; age 59-79 years, mean 68.0 ± 5.8), quantifying the tendency to experience complex and simple hallucinations in an experimental environment designed to elicit hallucinations. We find that reports of complex hallucinations (those representing objects, scenes, etc) are lower in older adults than in younger adults, both in real time and retrospectively. None of our measured cognitive or sensory measures (visual acuity, contrast sensitivity, perceptual reorganisation, imagery vividness, memory inhibition, and suggestibility) mediate this relationship. We conclude that reduced complex hallucination proneness appears normative in ageing, and that experiencing hallucinations in older individuals may signal underlying pathology.

## 1. Introduction

Visual hallucinations - percepts without corresponding external stimuli - are associated with various clinical conditions including schizophrenia (Silverstein & Lai, 2021), Parkinson’s disease (Barnes & David, 2001), dementia (Zarkali, Adams, et al., 2019), epilepsy (Kasper et al., 2010) and vision impairment (Jan & del Castillo, 2012). Many of these conditions are particularly prevalent in older adults. Though hallucinations are commonly experienced in these pathological states, they can also be experienced within the general population - both spontaneously (Johns et al., 2014; McGrath et al., 2015; Tien, 1991) and induced through experimental methods, for example using alterations in light and sound (Allefeld et al., 2011; Reeder, 2022; Wackermann et al., 2002). Across all of these states, the hallucinations that are experienced can range from simple (i.e. abstract, geometric forms) to more complex, (i.e. objects, human figures, or scenes) (Barnes & David, 2001; Jan & del Castillo, 2012; Kasper et al., 2010; Shenyan et al., 2024; Silverstein & Lai, 2021). However, it is unclear how susceptibility to phenomenologically distinct types of hallucinations changes across the lifespan in healthy adults.

Among the few studies that explore hallucination frequency in typical ageing, findings are inconsistent (Badcock et al., 2017, 2020a; Larøi et al., 2019; Shenyan et al., 2024; Soulas et al., 2016; Thompson et al., 2021). Though some studies report an increase in hallucinatory prevalence with age (Shenyan et al., 2024; Tien, 1991; Turvey et al., 2001), many report a decrease (Larøi et al., 2005, 2019; Maijer et al., 2018; Thompson et al., 2021). These studies often rely on self-report questionnaires, which may be prone to various biases: for example, older individuals may retrospectively under-report hallucinations due to stigma (Badcock et al., 2017, 2020a). We sought to resolve this discrepancy by measuring changes in hallucinatory proneness in a healthy ageing population using more objective, experimental measures (Shenyan et al., 2024). If hallucinations are rare in older people, their presence could provide a strong indicator of underlying pathological processes (Compton et al., 2012; Pagonabarraga et al., 2016; Zarkali, Adams, et al., 2019). Alternatively, an increase in hallucinations may simply be a normal aspect of ageing, and thus, we would observe a corresponding increase in a healthy, ageing population (Baumeister et al., 2017).

### 1.1 Age-related mechanisms of hallucinatory proneness

Ageing also offers a unique opportunity to test existing hypotheses about the multiple interrelated factors which have been shown to promote hallucinatory experiences. One key factor is an overreliance on prior expectations (Teufel et al., 2015; Zarkali, Adams, et al., 2019), which may act to increase the tendency to experience hallucinations when paired with an independent, pathological mechanism, for instance, in Lewy body disease (Zarkali, Adams, et al., 2019) or psychosis (Teufel et al., 2015). Alternatively, prior expectations may become overweighted to compensate for age-related declines in visual function, which reduce the reliability of sensory input (Collerton et al., 2023; Occelli et al., 2017). Given that visual function diminishes with age (Pitts, 1982), increased hallucinatory proneness in older adults may stem from the interaction between sensory deficits and stronger reliance on prior knowledge (Chan et al., 2021). We therefore predicted that both visual impairments and the weighting of prior expectations would increase with age, and that these factors would be associated with a heightened tendency to experience complex hallucinatory imagery in later life. Similarly, the strength and vividness of mental imagery - a predominantly top-down process (Dijkstra et al., 2017, 2019, 2022), may also influence hallucinatory proneness (Königsmark et al., 2021; Reeder, 2022). However, as mental imagery vividness tends to decline across the lifespan (Gulyás et al., 2022), this could alternatively predict a reduction in hallucinatory experiences in older age.

Another contributing factor to hallucinatory proneness is a reduced ability to suppress irrelevant signals—a cognitive process related to inhibitory control—which has been associated with increased auditory hallucinations (Alderson-Day et al., 2019; Badcock & Hugdahl, 2014; Swyer & Powers, 2020). This tendency can be measured through intentional memory inhibition tasks, which require participants to suppress responses to irrelevant memories (Alderson-Day et al., 2019; Paulik et al., 2007). Performance on tasks measuring inhibitory control and memory suppression correlate with the severity of hallucinations both in individuals with psychosis (Badcock et al., 2005; Waters et al., 2003) and in healthy controls, as measured via self-reported questionnaire (Alderson-Day et al., 2019). While most research on intentional memory inhibition and hallucinations has predominantly focused on the auditory domain, here we test the relationship between inhibition and visual hallucinations. Prior research suggests that the ability to intentionally suppress unwanted memories diminishes with age, with older adults’ performing poorer than younger adults in similar tasks (Anderson et al., 2011; Collette et al., 2009). This led us to hypothesise that age-related declines in intentional memory inhibition could also increase the tendency to experience hallucinatory experiences in older age.

Finally, individual differences in suggestibility may also play a role in hallucinatory proneness. For instance, hallucinatory-prone participants have been found to report voices in white noise more frequently when they are specifically instructed to expect the voice (Alganami et al., 2017). Additionally, various measures of trait suggestibility have been found to correlate with self-reported hallucinatory experiences (Alganami et al., 2017). This may be related to phenomenological control – the tendency for individuals to shape their perceptual experiences in response to suggestions or task demands (Dienes & Lush, 2023; Lush et al., 2020, 2021). Notably, suggestibility has also been observed to increase with age (Biondi et al., 2020; Mitchell et al., 2003; Page & Green, 2007), leading us to hypothesise that increases in this trait with age could contribute to increased hallucinatory experiences in older adults.

### 1.2 Current study

Two common ways of inducing visual hallucinatory imagery via experimental manipulations are through perceptual deprivation, i.e., presenting homogeneous visual and auditory inputs (Ganzfeld; Schmidt et al., 2020), and visual stimulation via a high frequency, salient flicker (Ganzflicker; Königsmark et al., 2021). Using these methods, we previously outlined a novel approach to measuring hallucinatory proneness, quantifying experiences in real-time using button presses and verbal descriptions, and retrospectively via drawings and questionnaires (Shenyan et al., 2024). These methodologies can consistently induce both ‘simple’ and ‘complex’ hallucinations, where simple hallucinations are abstract alterations in colour, shapes and patterns likely to be driven by bottom-up activation (Bressloff et al., 2001, 2002; Ermentrout & Cowan, 1979), and complex hallucinations are figurative percepts with relatively greater semantic associations, such as objects, faces and scenes, likely derived from a comparatively more top-down driven mechanism (Allefeld et al., 2011; Reeder, 2022; Shenyan et al., 2024).

We reasoned that by putting participants in an abstract, experimental situation where they might even be expected to hallucinate, we would mitigate stigma and anxieties around the associated implications of hallucinating which might occur during self-report of hallucinations in day-to-day contexts. Our hypotheses and predictions are primarily tailored towards the tendency to experience complex imagery, as complex hallucinations are less affected by strictly bottom-up processes (i.e. the intensity of visual drive), and therefore are more likely to be affected by risk factors related to higher-order cognitive processes, such as inhibition and an overreliance on prior expectations. Furthermore, complex hallucinations are likely more similar to the pathological hallucinations experienced in conditions like Parkinson’s Disease and schizophrenia (Barnes & David, 2001; Silverstein & Lai, 2021), and their prevalence in a healthy population may therefore have more relevance to clinical decision making.

We therefore conducted a comprehensive investigation into the tendency to experience experimentally-induced hallucinations in older adults, with a particular focus on whether the frequency and intensity of complex hallucinatory imagery varies with age. Building on findings from our previous study in a younger adult sample (Shenyan et al., 2024), and considering that numerous risk factors for hallucination proneness increase with age (as detailed above), we investigated how specific cognitive and perceptual factors hypothesised to influence hallucinatory experiences change from early to late adulthood. We further explored how these age-related changes might modulate the propensity to experience hallucinations. The factors of interest included reduced quality of sensory input, increased contribution of knowledge in perception, vividness of mental imagery, inhibitory control and suggestibility. We experimentally tested their relationship with the tendency to experience complex hallucinations during both a Ganzfeld and a Ganzflicker, as well as their association with age.

Our primary hypothesis was that both the number and intensity of complex hallucinations (those related to objects, faces, scenes, etc) would increase with age, related to age-related declines in visual function, increases in weighting of prior expectations, reduced inhibitory control, and increased suggestibility. Alternatively, it is also plausible that the frequency and intensity of complex hallucinations decreases with age, given evidence for a decline in the vividness of visual imagery across the lifespan.

To quantitatively assess these hypotheses, we collected several measures. Visual function was assessed through visual acuity and contrast sensitivity. Reliance on prior knowledge in perception was measured using a two-tone image task, wherein participants initially view an ambiguous, difficult to parse image. After being shown the original undistorted image, they view the ambiguous version again; improved recognition on this second viewing reflects the influence of prior knowledge on perception. Inhibitory control was assessed via the Inhibition of Currently Irrelevant Memories (ICIM) task, which evaluates participants’ ability to intentionally suppress memory-based interference during a sequential image recognition task using black-and-white line drawings. The vividness of visual imagery and suggestibility were measured using validated self-report questionnaires.

As an exploratory facet to our study, we also examined the prevalence of simple hallucinations (those related to more abstract patterns and forms) with age. Although we had no specific a priori hypotheses, the tendency to experience simple hallucinations is linked to cortical excitability (Bressloff et al., 2002; DaSilva Morgan et al., 2022; Shenyan et al., 2024). Since cortical *hypo*excitability has been observed with ageing (Cespón et al., 2022; Clark & Taylor, 2011) it is possible that simple hallucinations may decrease with age. Conversely, vision loss in healthy ageing could be paired with visual phenomena (i.e. floaters, phosphenes; (Bergstrom & Czyz, 2025) in a way that could be interpreted as simple hallucinations, leading to an increase in simple hallucinations.

Our results did not correspond to our prediction that reduced vision would lead to an increased reliance on prior knowledge and therefore an increased tendency to experience complex hallucinations. Instead, we found that complex hallucinations are reduced in old age. In our sample, we also found evidence for reduced suggestibility in old age, which could in principle explain reduced experiences of complex hallucinations. However, a mediation analysis provided no evidence for a causal relationship between complex hallucinations, age and suggestibility. These findings reveal that reduced visual ability, as occurs in healthy ageing, does not increase the likelihood of experiencing hallucinations. Our findings also suggest that the tendency to experience complex hallucinations is not a normal part of the ageing process, and that their presence is therefore likely to be indicative of a pathological process.

## 2. Methods

### 2.1 Sample

There were 48 participants in the study, with four excluded. Two of these were due to technical difficulties during the Ganzflicker task, one due to prematurely terminating the Ganzflicker task due to discomfort, and one due to general lack of comprehension of the task during the training phase of the study. This left 44 participants (21 male, 23 female, mean age: 47.6 years, SD: 21.34, age range: 19-79). Of these, 22 participants were in the younger age group (14 female, 8 male, mean age: 27.2 years, SD: 5.54, age range: 19-39), and 22 were in the older age group (10 female, 12 male, mean age: 68 years, SD: 5.77, age range: 59-79).

Exclusion criteria included any history of seizure or adverse experience to flashing or flickering lights, any first-degree relatives with a history of epilepsy, any neurological disorder including migraine with aura, and any visual disorder other than short and long sightedness and astigmatism. Participants were recruited through university recruitment systems, the National Institute for Health Research (NIHR) Joint Dementia Research database, and word of mouth. All participants provided informed consent. The experimental procedure was approved by the Research Ethics Committee at University College London (protocol number 24159/001). The research was carried out in accordance with the tenets of the Declaration of Helsinki.

### 2.2 Study design

After an introduction to the study, all participants began the experiment by completing a variety of baseline questionnaires and tasks. Participants were first asked to complete online versions of the Cardiff Anomalous Perceptions Scale (CAPS), the Vividness of Visual Imagery Questionnaire (VVIQ) and the Short Suggestibility Scale (SSS) questionnaires. Participants were then taken into a separate room where they underwent tests to assess their visual function and completed a two-tone task and an inhibitory control task, described in detail further down.

After a short break, participants began the Ganzeld and Ganzflicker procedures described in detail elsewhere (Shenyan et al., 2024) and below in the *Hallucination-inducing tasks* section (illustrated in Figure 1). Participants underwent 15 min of Ganzflicker and 25 min of Ganzfeld in a counterbalanced, repeated measures design. During an approximately thirty-minute break between conditions, participants drew their hallucinatory experiences with respect to their given descriptions and completed two retrospective questionnaires, the Altered States of Consciousness Rating Scale (ASC) (Dittrich et al., 2010) and the Imagery Experience Questionnaire (IEQ) (Roseby et al., 2020) to assess the intensity of their hallucinatory experiences.

**Figure 1:**
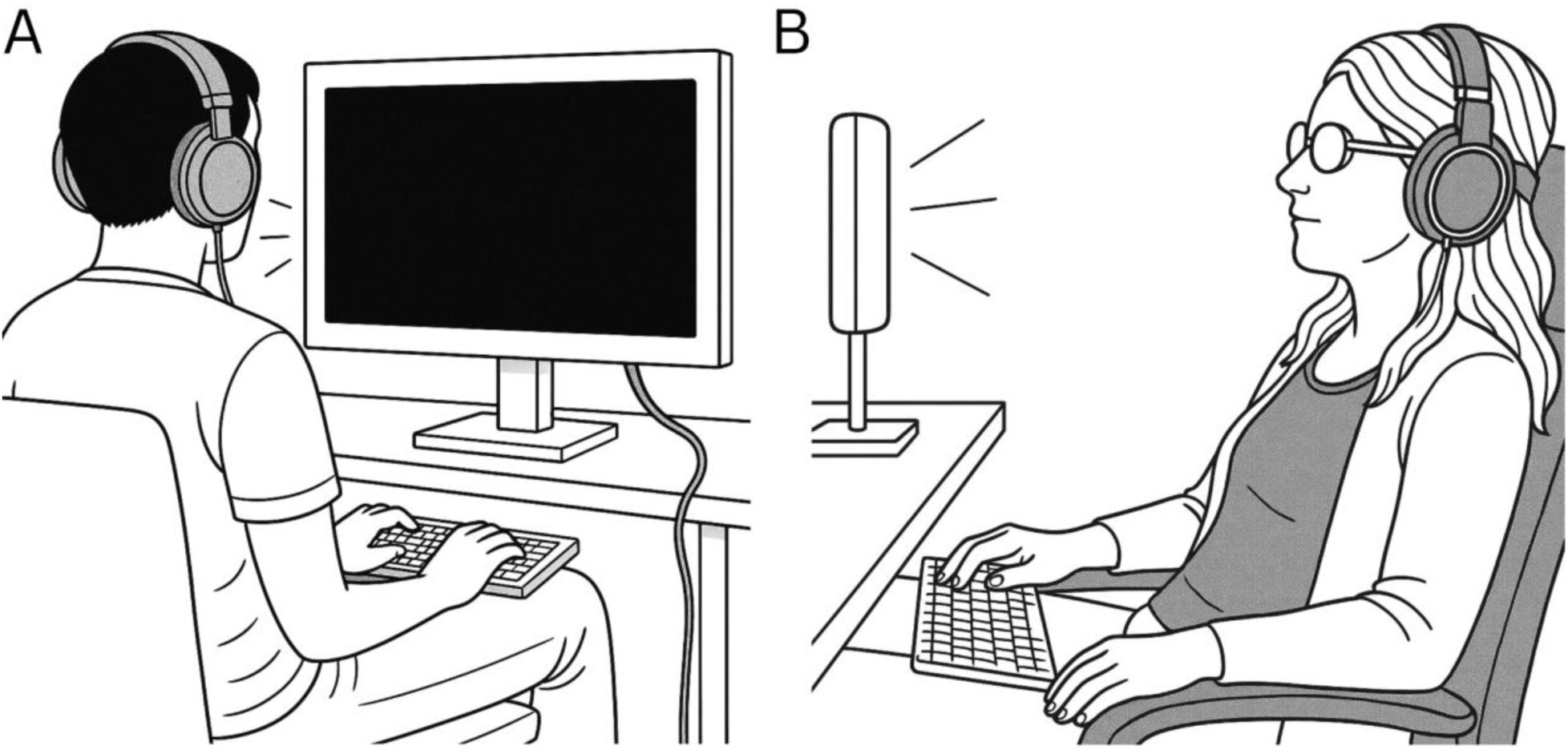
Schematic of participants during hallucination-inducing experimental techniques. In both conditions, participants are wearing headphones playing auditory brown noise with fingers placed on the space-bar of a keyboard to indicate the onset and offset of hallucinations. (A) Ganzflicker. Participants view a rapidly flickering red and black monitor (B) Ganzfeld. Participants are placed in front of a bright LED light, with ping pong ball halves placed over the eyes securely. Images are AI anonymoused illustrations of real photos depicting our testing set-up.

### 2.3 Experimental procedures

#### 2.3.1 Hallucination-inducing tasks

The instructions and procedures for the Ganzfeld and Ganzflicker tasks are fully described elsewhere (Shenyan et al., 2024). We provide a less detailed account in what follows.

##### 2.3.1.1 Instructions

Participants were given examples of the percepts that could arise: simple hallucinations such as colour, shapes, movements or patterns; complex hallucinations such as objects, faces and scenes; or nothing at all. Participants were encouraged to be patient, to keep an open mind, and to avoid active day-dreaming or mind wandering. Participants were asked to keep their eyes open, monitor any phenomena that arose across their visual field and to only verbalise a brief description of them once the percept had faded. These instructions were provided to ensure that participants reported all visual phenomena that could arise, even those that may have been more minor (i.e., simple, phosphene-like percepts) than they were expecting. In both procedures, auditory brown noise was played through noise-cancelling headphones and adjusted per participant to a volume that was comfortable, but loud enough to block out any external sound.

##### 2.3.1.2 Ganzflicker

Stimuli were coded using Psychtoolbox-3 running on MATLAB 2021b. Participants were seated in a darkened and soundproofed room, at ∼70 cm from a 32″ LED backlit LCD monitor (60 Hz frame rate; Cambridge Research Systems BOLDscreen) in a darkened and soundproofed room. The screen flickered an alternating black (luminance 0.38 cd/m2) and red (69.93 cd/m2) display for 15 min at a frequency of 10 Hz. Participants were trained on using the button press to indicate hallucinatory experience by undergoing a 30 s practice run during which they saw the flickering stimuli and practised pressing the space bar and verbalising their experience to the experimenter.

##### 2.3.1.3 Ganzfeld

Orange-coloured ping-pong balls were halved and taped securely over the eyes using medical tape. The visual field was illuminated by a warm white light (Lumary 24W Smart LED Flood Light). Participants were seated on a reclining chair approximately 70 cm from the light and were allowed to adjust their recline so that they were comfortable and relaxed. The position of the light was adjusted accordingly.

##### 2.3.1.4 Button press measures, drawings, and classification of hallucinations

Button-presses and verbal descriptions were used to describe the phenomenology of the hallucinations experienced by participants. The space bar of a keyboard was used to denote the onset of a hallucination. The space bar remained pressed until any hallucinatory form had faded, at which point, participants gave a brief verbal prompt of what they had seen. After the experiment, participants were given these descriptions to draw impressions of their hallucinatory experiences on an iPad (9th generation). These drawings, combined with the associated descriptions, gave the experimenter an idea of the nature of the hallucinations experienced and informed their classification as ‘simple’ or ‘complex’. Two independent experimenters scored hallucination complexity and met to discuss scoring discrepancies. Simple hallucinations were defined as any descriptions and corresponding drawings of colours, shapes, or patterns, while complex hallucinations were defined as drawings with corresponding semantic value and associated appropriate descriptions. When participants felt unable to draw or describe their hallucinations, they were classified as simple hallucinations.

##### 2.3.1.5 Post-questionnaires

Two abridged questionnaires were used to retrospectively assess the subjective experience of participants. We used the Altered States of Consciousness Rating Scale (ASC-R) (Dittrich et al., 2010), a well-validated 94-item self-report scale for the retrospective assessment of pharmacological and non-pharmacological induced altered states of consciousness. We chose questions primarily from the Elementary Imagery and Complex Imagery dimensions in line with our research question to interrogate the nature of participants’ subjective experience (i.e., questions from the *Simple Imagery* and *Complex Imagery*) subscales. The relevant items used within the study are in Supplementary Table S1.

We also utilised questions from the Imagery Experience Questionnaire (IEQ) (Roseby et al., 2020; Shenyan et al., 2024) to better capture the subjective visual experience of hallucinatory experiences. In line with our research question, we carried out analyses pertaining to hallucination complexity by separating items from the Complexity dimension of the IEQ into Simple (Items 1-4) and Complex (Items 5-8). The relevant items used within the study are in Supplementary Table S2.

#### 2.3.2 Behavioural tasks

##### 2.3.2.1 Two-tone task

###### 2.3.2.1.1 Behavioural stimuli generation

Stimuli were created using methods previously described elsewhere (Milne et al., 2022). In brief, 24 two-tone images were created through smoothing and binarising grayscale photographs of objects and animals in Matlab R2022a. Four ‘easy’ (without smoothing) two-tone images were used in practice trials and two were used as catch trials in the main experiment. All images were displayed at a fixed size of 13 degrees of visual angle on a mid-grey background.

###### 2.3.2.1.2 Procedure

Participants sat ∼70 cm in front of a 32″ (81cm) LED backlit LCD monitor (60 Hz frame rate; Cambridge Research Systems BOLDscreen) in a darkened and soundproofed room with the experimenter present. Participants first completed a training task in which a grayscale image was displayed and transformed gradually into a two-tone image that was unsmoothed and easy to recognise, to ensure they understood the relationship between two-tones and greyscale images. Next, participants completed four practice trials with unsmoothed ‘easy’ two-tone images. This was followed by the main task, in which 20 two-tone images of varying difficulty and the corresponding 20 greyscale images were presented in a pseudo-randomised order, such that for each grayscale image the corresponding two-tone was presented twice - once before the grayscale image (naive condition), and once after the grayscale image (cued condition). The task consisted of 60 experimental trials, with each trial consisting of a fixation cross presented for one second, followed by either a naive, greyscale or cued image presented for 200 ms, and finally a response screen. During the response screen, the question ‘Can you name what you saw?’ was displayed, prompting the participant to verbally name what they perceived in the image. The given answer was then typed in by the experimenter and displayed on the screen. Motivational, but uninformative, feedback was given after each trial to all participants. A diagrammatic of the task is illustrated in Figure 2. A break screen was displayed every 12 trials to show progress through the task. The task lasted approximately 15 minutes in total.

**Figure 2:**
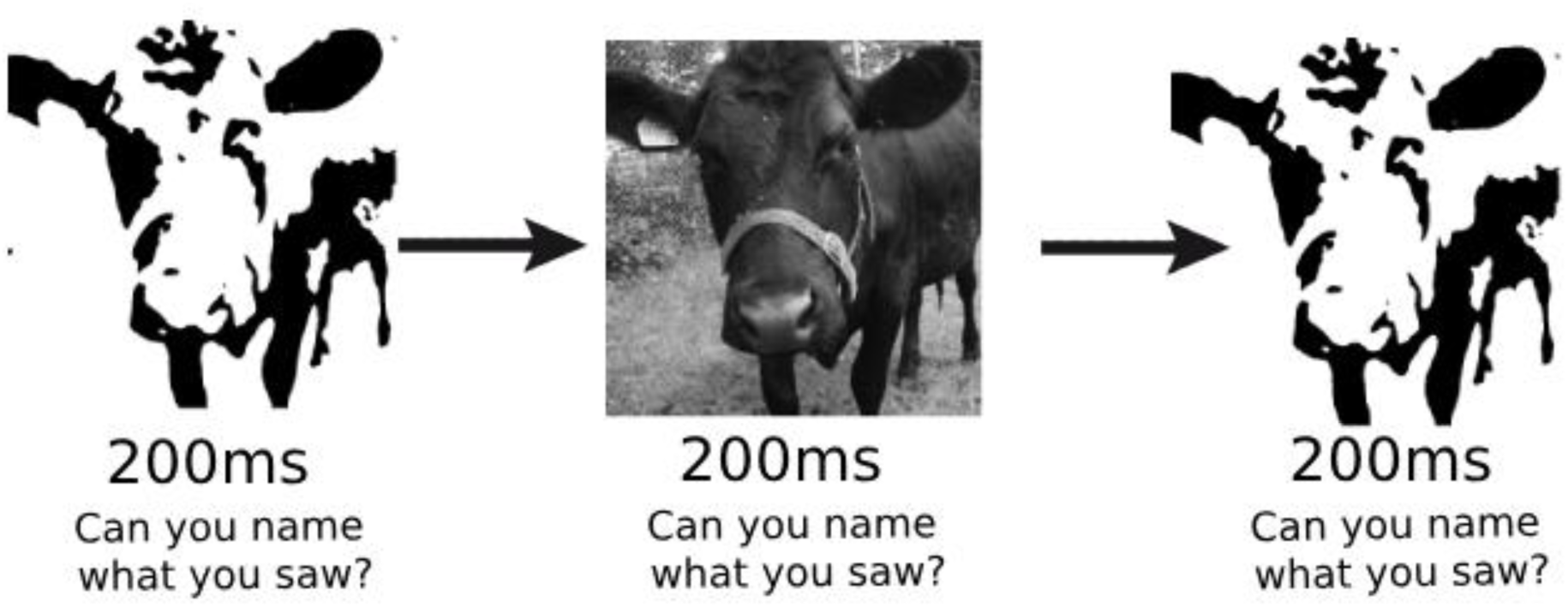
Diagrammatic illustration of the two-tone task utilised in the current experiment, which explores how prior knowledge is used to inform current perception. Participants were shown 60 experimental trials, with each trial consisting of a fixation cross presented for one second, followed by either a naive (not previously seen two-tone), greyscale or cued (two-tone following grayscale image presentation) image presented for 200 ms. After the image was presented for 200ms, a response screen was presented. During the response screen, the question ‘Can you name what you saw?’ was displayed, prompting the participant to verbally name what they perceived in the image. The given answer was then typed in by the experimenter and displayed on the screen.

###### 2.3.2.1.3 Scoring of behavioural performance

To quantify image recognition, we measured naive (first view of two-tone), greyscale and cued (view of two-tone after greyscale) naming accuracy. Image names were scored as correct if the content was correctly identified at the basic category level - superordinate categories names (e.g., naming a ‘cow’ an ‘animal’) were scored as incorrect; however, simple basic level or subordinate categories were accepted (e.g., a ‘tiger’ named as ‘cat’, or ‘scissors’ named as ‘shears’) as long as there was consistency in naming across the two-tone and greyscale conditions. For each participant, images that were not accurately identified in the greyscale condition were excluded from their data across all conditions. Two experimenters independently scored each image and met to resolve any discrepancies between scoring. Our primary outcome measure, the perceptual reorganisation effect, was quantified as the difference between the number of cued correct images and naive correct images, excluding trials where the grayscale image was named incorrectly, i.e.:

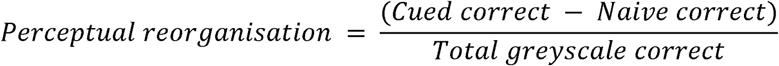

##### 2.3.2.2 The Inhibition of Current Irrelevant Memories task (ICIM)

The ICIM task was used to measure participants’ intentional memory inhibition ability through assessing image recognition in three blocks of sequential black-and-white line drawings (Alderson-Day et al., 2019). In each block, participants identified whether each image was a first presentation or a repeat *within the current block* using button presses (press “1” for first, press “2” for repeat within the block). This means that in the second and third blocks, participants had to distinguish whether a repeat was completely novel (press “1”), presented in a previous block (press “1”, inhibiting an irrelevant memory of seeing the object before), or presented before in **the current block** (press “2”). Per block, there were 95 image presentations, allowing for 35 possibilities to correctly identify a repeated image (i.e., “hit”) and 60 possibilities to incorrectly classify a first presentation as a repeat (i.e., “false alarm”) in each block. We measured task performance as the sum of false alarms from blocks two and three, as in previous literature (Alderson-Day et al., 2019).

#### 2.3.3 Visual function

Visual acuity and contrast sensitivity were measured to quantify participants’ visual functioning.

##### 2.3.3.1 Contrast sensitivity

Contrast sensitivity was assessed via a binocular Mars letter contrast sensitivity test (Mars Perceptrix, Chappaqua, NY; http://www.marsperceptrix.com/). This is a portable chart with eight rows of letters and six letters of constant size in each row (48 letters in total), which decrease in contrast at a rate of 0.04 log unit steps. The test was administered at 50 cm and illuminated by a lamp. Tests were scored with a value of 0.04 log contrast sensitivity per letter named correctly, with a maximum possible score of 1.92 log CS.

##### 2.3.3.2 Visual acuity

Visual acuity was assessed via a Good-Lite mini Snellen chart (Good-Lite, Elgin, IL; https://good-lite.com/); a compact, portable version of a traditional Snellen chart. The test was administered at 40 cm and illuminated by a lamp. Eye correction was measured in LogMAR units, ranging from 1.4 (worst) to -0.1 (best). Participants were asked to wear any eye correction worn on a day-to-day basis while completing the test.

#### 2.3.4 Pre-questionnaires

##### 2.3.4.1 CAPS

The Cardiff Anomalous Perceptions Scale (CAPS) is a 32-item measure of perceptual anomalies, such as hallucinations, in day-to-day life (Bell et al., 2006). We chose the CAPS as a secondary outcome measure of hallucinatory proneness, in order to assess whether day-to-day hallucinatory proneness, outside of an experimental context, also changes with age. Each item in the CAPS questionnaire is scored via a Yes (1) or No (0), and the total outcome measure comprises the sum of all the Yes values, with a maximum score of 32. Though the CAPS questionnaire also includes 3 separate subscales for the frequency, intrusiveness and distress associated with the perceptual experiences, these were not incorporated in the analyses for simplicity.

##### 2.3.4.2 SSS

The Short Suggestibility Scale (SSS) is a shortened form of the Multidimensional Iowa Suggestibility Scale (MISS), and assesses one’s susceptibility to accept and internalise external influences (Kotov et al., 2004). The scale consists of 21 items that are divided into categories of suggestibility, persuadability, sensation contagion, physiological reactivity and peer conformity. Each item is rated from 1 (“Not at all or very slightly”) to 5 (“A lot”), meaning the total outcome measure ranges from 21 to 105.

##### 2.3.4.3 VVIQ

The Vividness of Visual Imagery Questionnaire (VVIQ) assesses one’s mental imagery vividness(Marks, 1973). The questionnaire asks participants to form mental images of 16 scenes and scenarios, and to rate their vividness on a 5-point scale from 1 to 5. Therefore, the total outcome measure ranges from 16 to 80.

### 2.4 Statistical analysis

Our primary analyses involved either T-tests or Mann-Whitney U tests (depending on the normality of the data) to evaluate differences in the number of complex hallucinations pooled from both Ganzfeld and Ganzflicker conditions, between the older and younger age groups. We validated these findings by assessing differences in the Complex imagery sub-components in both the ASC and IEQ questionnaires. In order to determine whether day-to-day anomalous perceptual experiences outside of an experimental context varied between age groups, we compared CAPS scores between older and younger age groups.

To assess the factors which may affect age-related changes in hallucination complexity, we conducted Spearman’s rank correlations between the number of complex hallucinations and CS, VA, perceptual reorganisation score in the two-tone task, VVIQ scores, ICIM false alarm rates and SSS scores across our entire sample (N=44).

Where there was a difference in outcomes between groups as well as a statistically significant positive correlation between said outcome and the number of complex hallucinations, we conducted Bayesian mediation analyses. Models were estimated using the *brms*(Bürkner, 2017) package in R. Each mediation analysis involved two models: a mediator model predicting the mediator from age with a Gaussian family, and an outcome model predicting the outcome from both age and the mediator with a negative binomial family. As the outcome model is nonlinear, mediation effects are not constant but vary depending on the specific value of the predictor. Therefore, mediation was estimated by simulating counterfactual outcomes under two specific values of age: 27.2 and 68 years, corresponding to the average age of the younger and older groups in our sample (Pearl, 2012). We therefore computed, for each mediation analysis, the effects for a 40.2 year difference. The effects derived were the Total Effect (TE); the total effect of the age and the mediator variable on complex hallucinations; the Indirect Effect (IE); the portion of the total effect mediated by the change in the mediator, and the Direct Effect (DE), the effect of changing age from 27.2 to 68 while holding the mediator fixed at its value for age = 27.2. The posterior summaries of these effects were obtained using the bayestestR package, from which we report the mean and the 95% highest posterior density interval (HDI).

## 3. Results

We first present illustrative examples of complex and simple hallucinations drawn by our older and younger age participants in Fig 3. Associated abbreviated descriptions provided by participants accompany these drawings.

**Figure 3:**
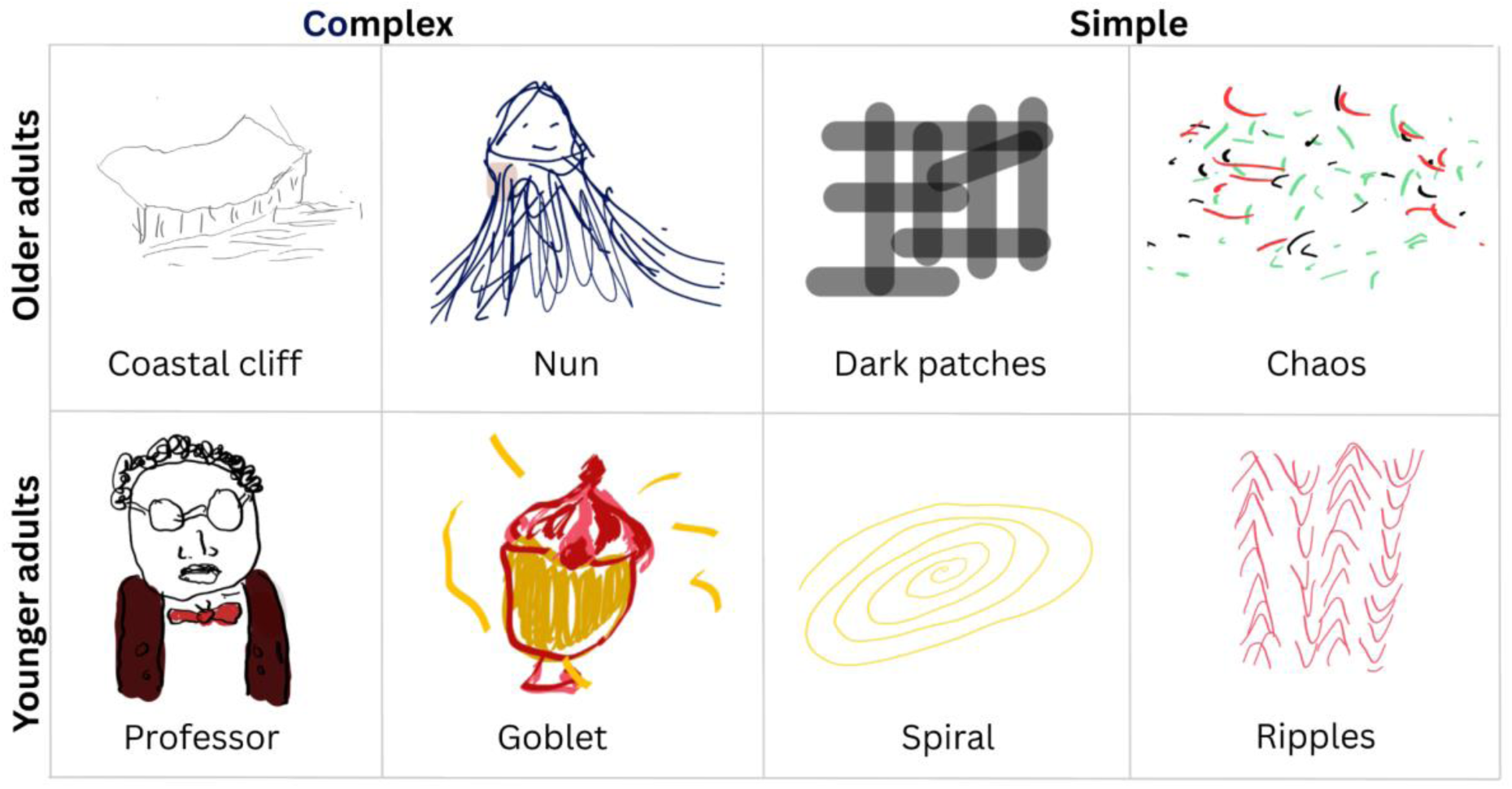
Example drawings of complex and simple hallucinations experienced during Ganzflicker and Ganzfeld in older adults and younger adults, accompanied with abbreviated descriptions given by participants.

### 3.1 Complex hallucination proneness in older-age and younger-age populations

During our hallucination-inducing protocols, the tendency to experience complex hallucinations was decreased in older adults across all outcome measures, contrary to our primary hypothesis. Button-press responses revealed that older adults reported fewer complex hallucinations than younger adults (Fig. 4A; W = 355.50, p = .006). Two separate, but correlated(Shenyan et al., 2024) questionnaires assessing the retrospective intensity of complex imagery during our experiments showed a similar numerical pattern, though only one of these questionnaires reached the threshold for statistical significance (IEQ - Fig. 4B; W = 351.50, p = .0097; ASC - Fig. 4C; W = 299, p = .122). Finally, scores on the CAPS questionnaire—a measure of abnormal perceptual experiences in daily life—were also lower in older adults compared to younger adults (Fig 4D; t = 3.08, p = .004).

**Figure 4:**
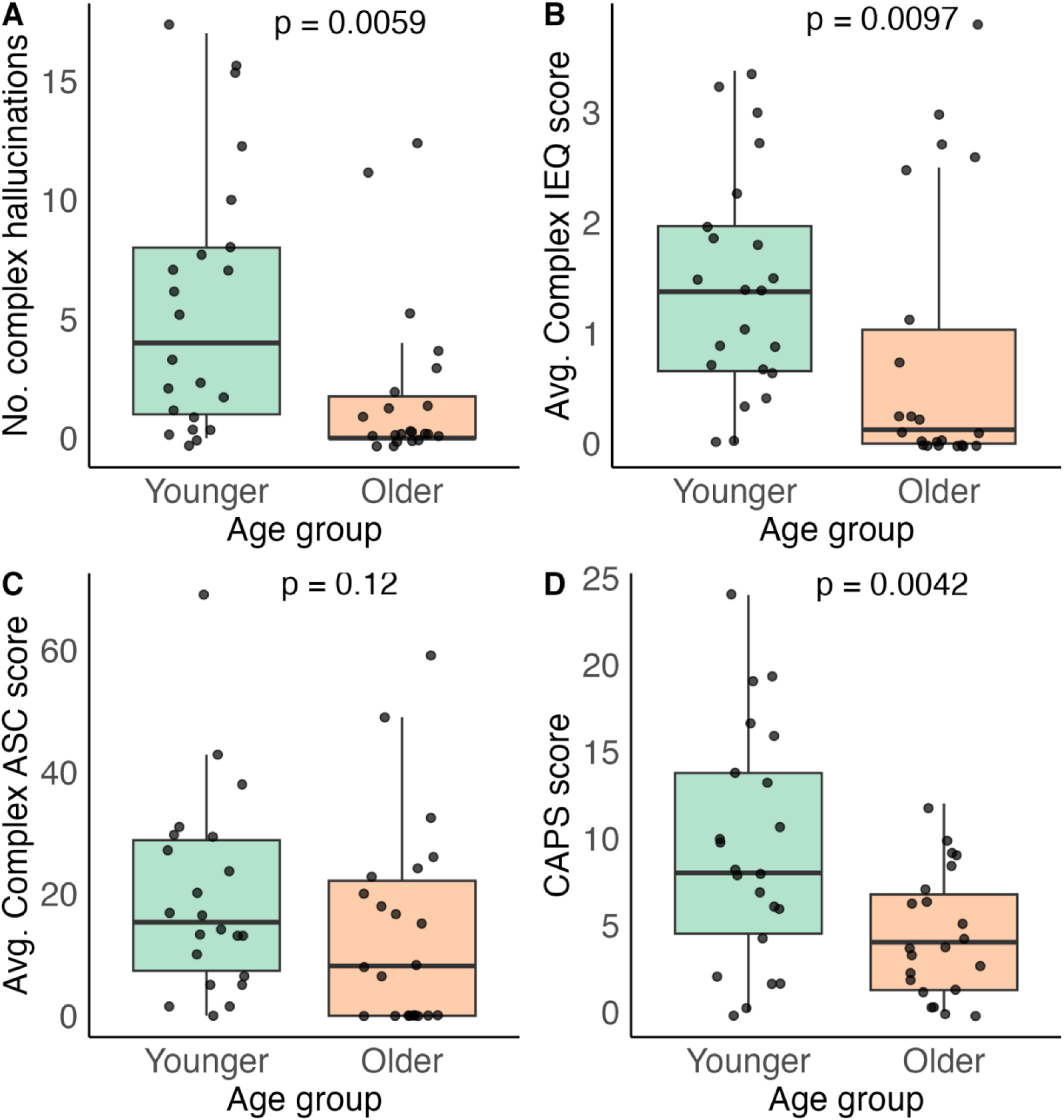
Differences in various measures of complex hallucinations during Ganzfeld and Ganzflicker between younger adults (green) and older adults (orange) (x-axis) (A) Numbers of complex hallucinations (y-axis) (B) Average retrospectively rated intensity of complex hallucinatory imagery, as measured through the Imagery Experience Questionnaire (IEQ) (y-axis) (C) Average retrospectively rated intensity of complex hallucinatory imagery, as measured through the Altered States of Consciousness Rating Scale (ASC) (y-axis) (D) Day-to-day anomalous perceptual experiences (y-axis), as measured through the Cardiff Anomalous Perceptual Scale (CAPS)

### 3.2 Simple hallucination proneness in older-age and younger-age populations

We observed no differences in the number of simple hallucinations between younger adults and older adults reported via button press (Figure 5A; W=221, p=0.63). Both questionnaires used showed a numerical decrease in the retrospective intensity of simple hallucinatory experiences, though only one of these questionnaires reached the threshold for statistical significance (IEQ - Figure 5B; W = 376.5, p = .0016; ASC - Figure 5C; W = 319.5, p = .071). On the whole our results suggest that the rate of simple hallucinations shows little-to-no change in the course of healthy ageing, particularly in comparison to the rate of complex hallucinations – though there may be some changes in the intensity of simple hallucinations.

**Figure 5:**
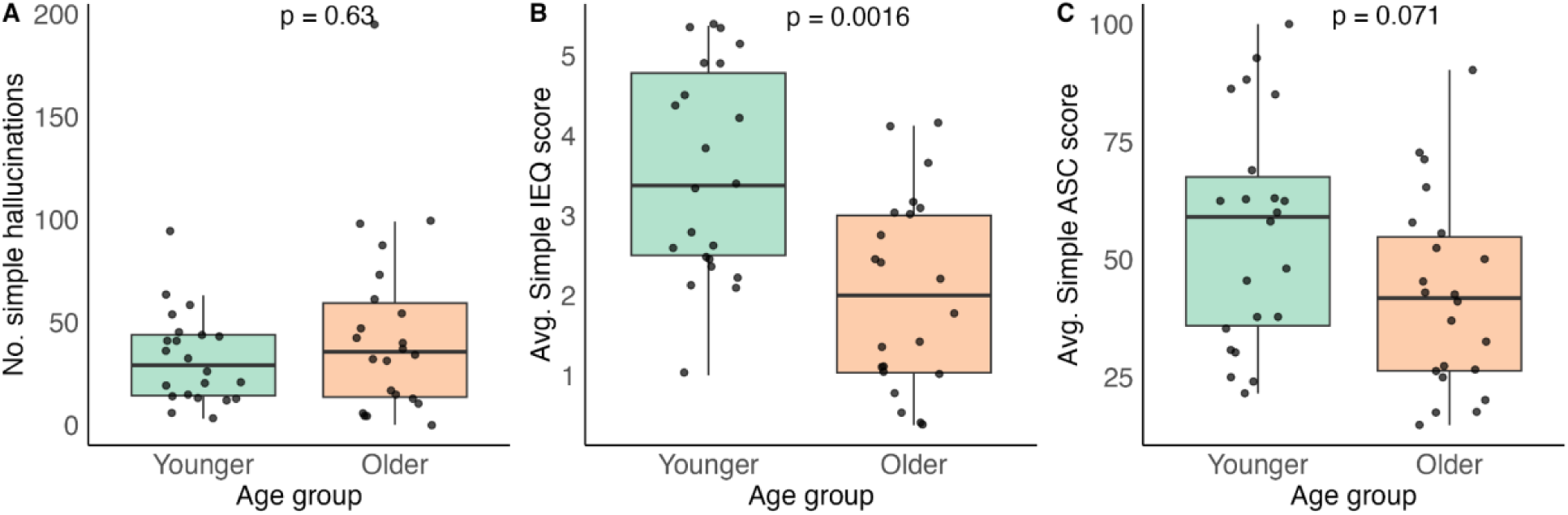
Differences in various measures of simple hallucinations during Ganzfeld and Ganzflicker between younger adults (green) and older adults (orange) (x-axis) (A) Numbers of simple hallucinations (y-axis) (B) Average retrospectively rated intensity of simple hallucinatory imagery, as measured through the Imagery Experience Questionnaire (IEQ) (y-axis) (C) Average retrospectively rated intensity of simple hallucinatory imagery, as measured through the Altered States of Consciousness (Rating Scale) ASC (y-axis).

### 3.3 Reduced sensory input and up-weighted priors

Having established a reduction in hallucination complexity in older-age adults, we next explored individual factors that could explain this change. We first considered whether deficits in bottom-up input (i.e., changes in contrast sensitivity and visual acuity) differed between our older and younger age groups, whether these measures were correlated with the number of complex hallucinations, and finally, whether these measures mediated the relationship between age and the number of complex hallucinations.

Older adults had significantly worse visual acuity and contrast sensitivity than younger adults (visual acuity - Fig 6A; W = 24, p < .001; contrast sensitivity - Fig 6B; W = 406.5, p < .001). The correlation between contrast sensitivity and complex hallucinations was non-significant (Fig 6F; rs(42) = .14, p = .36). Though visual acuity scores were negatively correlated with the number of complex hallucinations (Fig 6E; rs(42) = -.34, p = .026), this was no longer significant after controlling for age (rs(42) = .07, p = .67). There were also no significant correlations of interest between numbers of simple hallucinations and either of these measures (contrast sensitivity - rs(42) = .16, p = .3; visual acuity - rs(42) = .01, p = .94).

**Figure 6:**
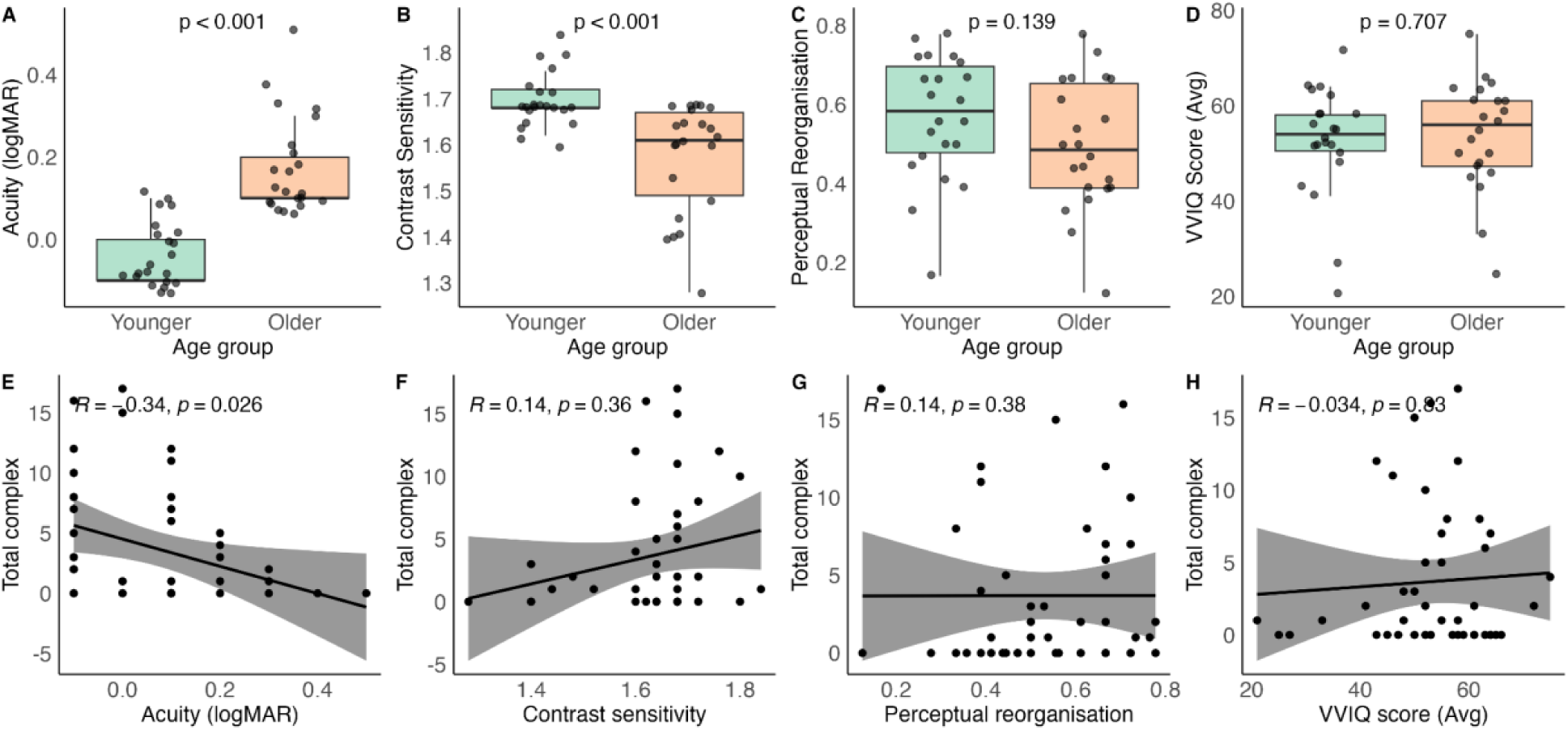
Relationship between age, complex hallucinations, and bottom-up (visual acuity, contrast sensitivity) and top-down (perceptual reorganisation, Vividness of Visual Imagery (VVIQ) questionnaire scores) measures. Difference in younger (green) and older adults (orange; A-D) and scatter plots showing relationship to complexity (E-H). (A) Visual acuity in logMAR units (y-axis), where higher logMAR acuity is indicative of worse vision; (B) Contrast sensitivity (y-axis); (C) Perceptual reorganisation scores on two-tone task (D) VVIQ scores. Scatter plots showing relationship between total number of (E) complex hallucinations (y-axis) and visual acuity (x axis) with associated correlation coefficient; (F) complex hallucinations (y-axis) and contrast sensitivity (x axis); (G) complex hallucinations (y-axis) and perceptual reorganisation score on two-tone task (x axis); and (H) complex hallucinations (y-axis) and average VVIQ score (x axis). All correlations shown are Spearman’s rank correlation tests with associated black trendline and 95% CI grey shading.

We then ran non-linear Bayesian mediation analyses to ascertain whether decreases in visual acuity mediated the relationship between age and decreased complex hallucinations. Posterior estimates suggested a Total Effect of the model, i.e., the overall total impact of age and the mediator, visual acuity, on complex hallucinations (*M* = –4.93, 95% CI [–9.88, –0.9]), with strong evidence of a non-zero effect (pd = 99.72%). The direct effect of age, independent of visual acuity, was also likely non-zero (*M* = –4.70, 95% CI [–10.35, -0.2], pd = 98.55%). The indirect effect of age mediated through visual acuity was small and uncertain (*M* = –0.23, 95% CI [– 1.67, 0.96]), pd = 57.67%). These results suggest that while age is associated with a decreased rate of complex hallucinations, visual acuity explains only a negligible part of this relationship, with the effect primarily operating through a direct pathway.

We then considered whether changes in the weighting of top-down processes - for instance, the degree of perceptual reorganisation in a two-tone task and vividness of mental imagery, as measured through the VVIQ, relate to hallucination complexity and age. As before, we first considered whether there were differences between these measures between our younger and older age groups, whether these measures were correlated with the number of complex hallucinations, and finally whether these measures mediated the relationship between age and the number of complex hallucinations.

Although mean perceptual reorganisation scores were numerically lower in older adults compared to younger adults, this was not statistically significant (Fig 6C; t = 1.51, p = .14), nor was there a significant correlation between perceptual reorganisation scores and complex hallucinations (Fig 6G; rs(42) = .14, p = .38). There was no statistically significant difference between the VVIQ scores of younger and older adults (Fig 5D; W = 227.5, p = .74) nor was there a significant correlation between VVIQ scores and complex hallucinations (Fig 6F; rs(42) = - .034, p = .83). As expected, there were no significant correlations between any of our top-down measures and the tendency to experience simple hallucinations (perceptual reorganisation – rs(42) = - .0051, p = .97; VVIQ – rs(42) = - .17, p = .26). As neither the difference between age groups in these measures nor the correlation between the number of complex hallucinations and these measures was significant, we did not conduct further mediation analyses.

### 3.4 Inhibition

There was no significant difference between ICIM false alarm rates between older adults and younger adults (t = 1.13, p = .27). There was also no significant relationship between ICIM false alarms and the number of complex (rs(42) = -.031, p = .85) or simple (rs(42) = -.27, p = .11). Again, we did not conduct further mediation analyses.

### 3.5 Suggestibility

Finally, we considered the exploratory relationship between suggestibility, complex hallucinations and age. Mean SSS scores were lower in older adults compared to younger adults (Fig 7A; t = 4.88, p < .001). There was also a significant relationship between suggestibility and the number of complex hallucinations (Fig 7B; rs(42) = .32, p = .032), however, this was no longer significant after controlling for age (rs(42) = .003, p = .98). There was no significant relationship between simple hallucinations and SSS scores (rs(42) = .19, p = .22).

**Figure 7:**
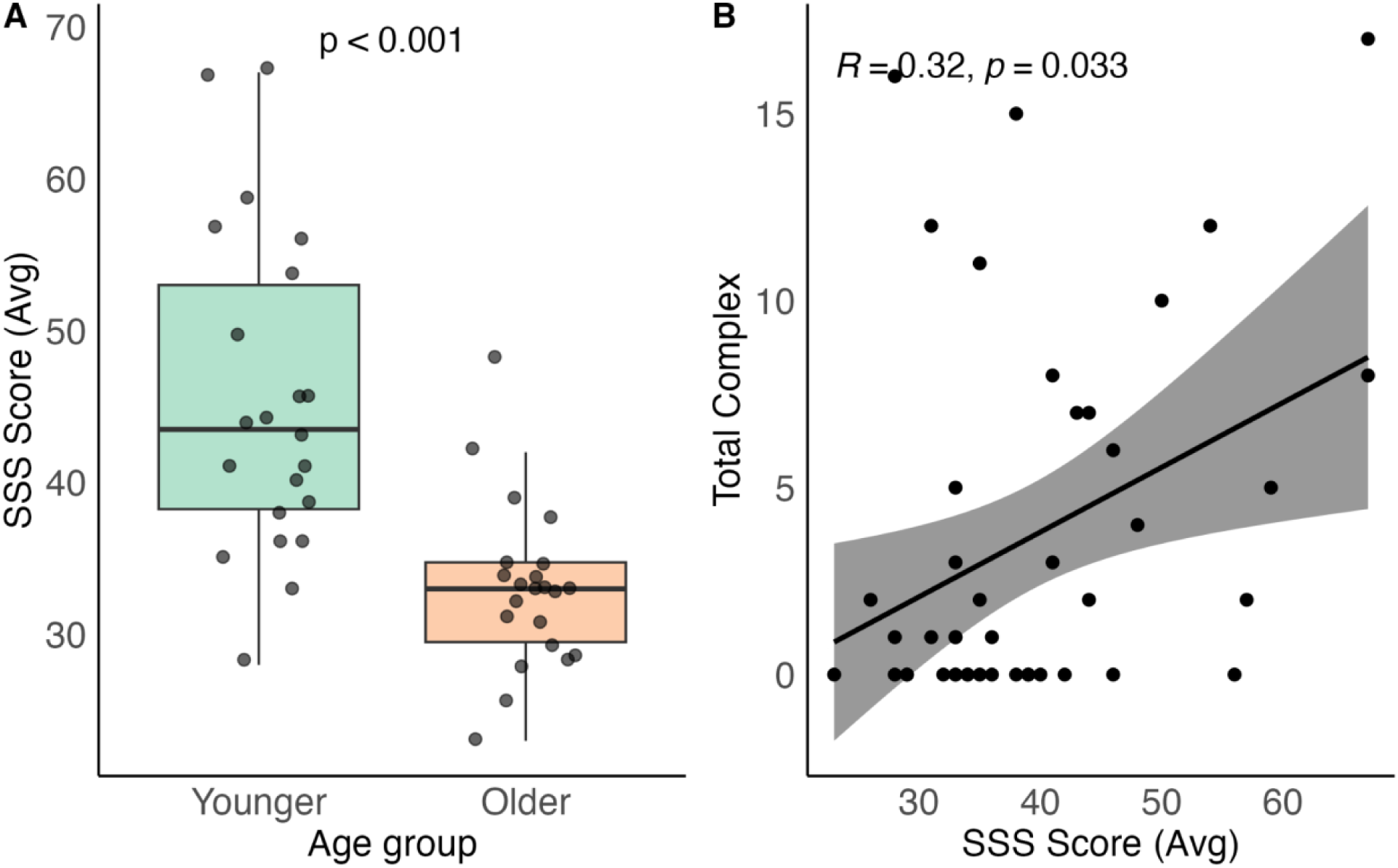
Relationship between age, complex hallucinations, and the Short Suggestibility Scale (SSS). (A) Differences in average SSS scores (y-axis) between younger adults (green) and older adults (orange) (x-axis) (B) Scatter plot showing relationship between total number of complex hallucinations (y-axis) and average SSS score (x axis). All correlations shown are Spearman’s rank correlation tests with associated black trendline and 95% CI grey shading.

We again carried out a Bayesian mediation analysis to see whether decreases in suggestibility in old-age explained decreases in numbers of complex hallucinations. We see a Total Effect of age and suggestibility combined on complex hallucinations (*M* = –4.93, 95% CI [–9.98, –0.9]), with very strong evidence for a non-zero effect (pd = 99.70%). The direct effect of age, independent of suggestibility, on complex hallucinations was also strongly supported (*M* = – 4.70, 95% CI [–10.12, 0.05], pd = 98.60%). The indirect effect via suggestibility was small and highly uncertain (*M* = –0.23, 95% CI [–1.70, 0.9], pd = 58.10%). These results suggest that while age is associated with decreased complex hallucinations, suggestibility does not appear to meaningfully mediate this relationship, with the effect primarily driven by a direct pathway.

## 4. Discussion

In this study, we sought to investigate how susceptibility to non-pathological visual hallucinations changes with age, and to identify the various factors that contribute to these changes. Some questionnaire studies have shown a reduced tendency to experience hallucinations with age in day to day life (Kråkvik et al., 2015; Maijer et al., 2018; Soulas et al., 2016; Thompson et al., 2021). Other conflicting studies, however, suggest an *increase* in hallucination proneness with age (Shenyan et al., 2024; Tien, 1991; Turvey et al., 2001), and moreover many factors for hallucination proneness increase with age (i.e. reductions in visual function and cognitive abilities (Badcock et al., 2020b; Chan et al., 2021; Collette et al., 2009; Owsley et al., 1983; Pitts, 1982; Wynn et al., 2020). We therefore reasoned that reports of reduced hallucination proneness may be due to the stigma associated with experiencing hallucinations and the implications, and we predicted that placing participants in an abstract, controlled experimental paradigm designed to elicit hallucinations (and therefore minimising stigma-related underreporting) would reveal an age-related increase in hallucinations.

Contrary to our hypothesis, we found that the tendency to experience experimentally-induced complex visual hallucinations in a laboratory environment decreases with age, and there was no consistent age difference in the tendency to experience simple hallucinations. This was despite replicating substantial, well-documented declines in visual acuity and contrast sensitivity with age in our sample (Owsley et al., 1983; Pitts, 1982) This suggests that age-related sight loss does not increase hallucination proneness, unlike more severe forms of sight loss (Jan & del Castillo, 2012). In addition, processes thought to involve top-down modulation of sensory brain regions and hypothesised to correlate with hallucination proneness, such as perceptual reorganisation (Teufel et al., 2013; Zarkali, Adams, et al., 2019), vividness of mental imagery (Königsmark et al., 2021; Reeder, 2022), and memory inhibition (Alderson-Day et al., 2019; Paulik et al., 2007), showed no significant differences between our groups of healthy younger and older adults.

Older participants also exhibited reduced suggestibility, which correlated with fewer complex hallucinations. However, this association may simply reflect the decline of suggestibility with age, as we found no evidence for a causal relationship with complex hallucinations. Despite this, this work still highlights a need to explore suggestibility as a potential factor in reporting during hallucination reporting experiments, for example by increasing responding to meet perceived expectations or more lenient criterion setting. To disentangle this, future studies could also include control ‘faux hallucination induction’ conditions in which true visual hallucinations are unlikely to occur, but participants are prompted to report specific anomalous perceptual experiences, and to explore the relationship between percepts reported in the control conditions and those reported in true hallucination-inducing conditions.

Our findings add to a body of literature exploring hallucinations in a healthy ageing population in various ways. Firstly, for the first time, we experimentally induced visual hallucinations in older-adults through two distinct methods: visual stimulation via a rapidly flickering screen (Ganzflicker) and sensory deprivation using a static, homogeneous light field paired with auditory brown noise (Ganzfeld). We show that despite the numerous risk factors for increased hallucination-proneness typically associated with aging, such as reductions in sensory abilities (Owsley et al., 1983; Pitts, 1982), older adults report fewer complex hallucinations during experimental conditions as well as self-reporting the retrospective intensity of this complex imagery as lower. There are several potential mechanisms to explain these changes.

The lower incidence of hallucinations in our older adult sample, and their lack of association with expected risk factors, may reflect the influence of sample selection and the underlying dynamics of pathology across the lifespan. For example, traits such as increased top-down influence on perception and inhibition of irrelevant representations, have been linked to conditions like Lewy body dementia (Zarkali, Adams, et al., 2019) or psychosis (Teufel et al., 2015; Soriano et al., 2009) and may signal early vulnerability to pathology in younger individuals (Haarsma et al., 2020; Paulik et al., 2007; Teufel et al., 2013). Those whose hallucination-related traits progressed into overt clinical conditions may be underrepresented in older healthy samples due to an earlier onset and diagnosis. Thus, the reduced hallucination proneness in our older sample may reflect a filtering effect over time. Our work suggests that when hallucinations do occur in older adults, they may warrant closer clinical attention, as they could signal underlying pathology rather than normative age-related change. Future research could use controlled hallucination-inducing paradigms, such as flicker or sensory deprivation, as safe “stress tests” (Zarkali, Lees, et al., 2019) to probe individual differences in hallucination susceptibility and potentially flag early signs of neurodegenerative or psychiatric disorders.

Age-related cortical hypoexcitability (Cespón et al., 2022; Clark & Taylor, 2011) could also explain the decline of complex hallucinations we observed in our older adult sample. Our focus here has been on top-down modulatory factors as the primary driver of complex hallucinations, given their relative insensitivity to bottom-up manipulations (Shenyan et al., 2024). However, these findings do not negate a contributory role for intrinsic cortical excitability in generating complex hallucinations. Simple hallucinations have been linked with cortical excitability (Bressloff et al., 2001, 2002), and though we observed no differences in the rate of button-press reports for simple hallucinations, we did see evidence to suggest a decrease in the retrospectively rated intensity of simple hallucinations in old age, which could also be driven by cortical hypoexcitability. Future studies should investigate this possibility, for example by using methods such as transcranial magnetic stimulation (TMS) to measure flicker phosphene thresholds in both older and younger adults (Abrahamyan et al., 2011).

## Limitations

As measures of hallucinations rely on self-reported measures, there is still the possibility that older individuals may conservatively under-report hallucinations due to stigma. Furthermore, our younger cohort was composed mostly of undergraduate university students. Therefore, there may also be a cohort effect, where younger adults are more accepting of the possibility of experiencing hallucinatory experiences during a psychology experiment. We further did not collect background data on socioeconomic factors like education levels, which could vary by age and influence hallucinatory reporting. These factors therefore act as potential unmeasured mediators, which may inflate the estimates of the direct effect of age on hallucination proneness in our mediation analyses.

Furthermore, the classification of hallucinations as complex in our study was based on researcher judgement, which inherently introduces potential bias. In order to mitigate this, we were particularly conservative in our classification of complex hallucinations – concordance between the provided description and the drawing was required to be appropriately classified as a complex hallucination. Moreover, discrepancies in classifications were discussed among the experimenter and other researchers involved in the project to reach a consensus. While these measures were implemented to reduce bias and enhance the reliability of our findings, it is important to acknowledge that some degree of researcher bias may still persist (Shenyan et al., 2024). In addition, the dynamic nature of hallucinations, specifically during the Ganzflicker (Shenyan et al., 2024), presents challenges for verbal reporting. It is possible that some participants may have experienced visual hallucinations but were unable to effectively articulate them. However, the consistency of our results both directly measured inside the laboratory (i.e., button press measures and self-report retrospective questionnaires) and outside of the laboratory (i.e., CAPS which measures anomalous perception in day-to-day life), decreases the possibility of both of these situations. Incorporating objective measures such as neuroimaging techniques could provide further insights into the neural differences between younger and older adults during these paradigms.

## Conclusions

To conclude, we show that older adults are less prone to experimentally-induced visual hallucinations than younger adults, aligning with a growing body of literature. Our findings suggest that when hallucinations do occur in older adults they may be indicative of underlying pathology, rather than being a result of the risk factors associated with normal ageing (i.e., sensory and cognitive impairments). Further research is needed to explore whether we are able to use safe hallucination-inducing environments - such as flicker or sensory deprivation - as ‘stress tests’ for hallucination-associated pathology, as well as cognitive factors which contribute to complex hallucination formation and how these differ across the lifespan.

## Data availability statement

Data is available on request. For data requests, please contact the corresponding author.

## Authorship contribution statement

OS: Conceptualisation, Methodology, Data collection, Analysis, Visualisation, Writing – Original draft preparation. LH: Data collection, Analysis, Writing – Reviewing and editing. GM: Methodology, Writing – Reviewing and editing. ML: Analysis, Writing – Reviewing and editing. JAG: Supervision, Writing – Reviewing and editing. JIS: Conceptualisation, Methodology, Supervision, Writing – Reviewing and editing. TMD: Conceptualisation, Methodology, Supervision, Writing – Reviewing and editing. All authors reviewed and approved the final manuscript.

## Declaration of generative AI and AI-assisted technologies in the writing process

During the preparation of this work the authors used ChatGPT in order to create AI anonymoused illustrations of real photos depicting testing set-ups (Figure 1). After using this tool, the authors reviewed and edited the content as needed and tak full responsibility for the content of the publication.

