## Supplementary materials for "Reduced susceptibility to experimentally-induced complex visual hallucinations with age"

### Supplementary methods

**Supplementary Table S1**: Items from ASC-R utilised in study

| Dimension | Item | Question |
| --- | --- | --- |
| Elementary Imagery (11D) | 14 | I saw regular patterns [in complete darkness or with closed eyes] |
|  | 22 | I saw colours before me [in total darkness or with closed eyes] |
| Complex Imagery (11D) | 39 | I saw scenes rolling by [in total darkness or with my eyes closed] |
|  | 72 | I could see pictures from my past or fantasy extremely clearly |
|  | 82 | My imagination was extremely vivid |

**Supplementary Table 2:** Items from IEQ utilised in study

| Dimension | Item | Question |
| --- | --- | --- |
| Complexity | 1 | I saw bursts of light or splashes of colour. |
|  | 2 | I saw abstract geometrical designs and patterns. |
|  | 3 | I saw rapidly transforming objects/ figures. |
|  | 4 | I saw repetitive, moving objects/ figures embedded in geometrical patterns. |
|  | 5 | I saw stable, well-defined objects/ figures. |
|  | 6 | I saw snapshots or glimpses of full scenes |
|  | 7 | I saw full-fledged scenes without being a part of them, similar to watching a movie |
|  | 8 | I was fully immersed within what looked and felt like another authentic realm |
